# Graded multidimensional clinical and radiological variation in patients with Alzheimer’s disease and posterior cortical atrophy

**DOI:** 10.1101/2023.02.07.527424

**Authors:** Ruth U. Ingram, Dilek Ocal, Ajay D. Halai, Gorana Pobric, David Cash, Sebastian J. Crutch, Keir X.X. Yong, Matthew A. Lambon Ralph

**Affiliations:** Division of Psychology and Mental Health, School of Health Sciences, University of Manchester, UK; Dementia Research Centre, UCL Institute of Neurology, London, UK; MRC Cognition & Brain Sciences Unit, University of Cambridge, UK

**Author notes:** **Corresponding authors,** Ruth U. Ingram, PhD, University of Manchester, Manchester, M13 9PL, Prof. Matthew A. Lambon Ralph, MRC Cognition & Brain Sciences Unit, University of Cambridge, UK. Joint first authors (Ruth U. Ingram, Dilek Ocal). Joint senior authors (Keir X.X. Yong, Matthew A. Lambon Ralph). **Open access**: For the purpose of open access, the UKRI-funded authors have applied a Creative Commons Attribution (CC BY) licence to any Author Accepted Manuscript version arising from this submission.

## Abstract

**Background and Objectives:** Alzheimer’s disease spans heterogeneous typical and atypical phenotypes. Posterior cortical atrophy is one striking example, characterised by prominent impairment in visual and other posterior functions in contrast to typical, amnestic Alzheimer’s disease. The primary study objective was to establish how the similarities and differences of cognition and brain volumes within Alzheimer’s disease and posterior cortical atrophy (and by extension other Alzheimer’s disease variants), can be conceptualised as systematic variations across a transdiagnostic, graded multidimensional space.

**Methods:** This was a cross-sectional, single-center, observational, cohort study performed at the National Hospital for Neurology & Neurosurgery, London, UK. Data were collected from a cohort of PCA and AD patients, matched for age, disease duration and MMSE scores. There were two sets of outcome measures: (1) scores on a neuropsychological battery containing 22 tests spanning visuoperceptual and visuospatial processing, episodic memory, language, executive functions, calculation, and visuospatial processing; and (2) measures extracted from high-resolution T1-weighted volumetric MRI scans. Principal component analysis was used to extract the transdiagnostic dimensions of phenotypical variation from the detailed neuropsychological data. Voxel-based morphometry was used to examine associations between the PCA-derived clinical phenotypes and the structural measures.

**Results:** We enrolled 93 PCA participants (mean: age = 59.9 yrs, MMSE = 21.2; 59/93 female) and 58 AD participants (mean: age = 57.1 yrs, MMSE = 19.7; 22/58 female). The principal component analysis for posterior cortical atrophy (sample adequacy confirmed: Kaiser-Meyer-Olkin = 0.865) extracted three dimensions accounting for 61.0% of variance in patients’ performance, reflecting general cognitive impairment, visuoperceptual deficits and visuospatial impairments. Plotting Alzheimer’s disease cases into the posterior cortical atrophy-derived multidimensional space, and vice versa, revealed graded, overlapping variations between cases along these dimensions, with no evidence for categorical-like patient clustering. Likewise, the relationship between brain volumes and scores on the extracted dimensions was overlapping for posterior cortical atrophy and Alzheimer’s disease cases.

**Discussion:** These results provide evidence supporting a reconceptualization of clinical and radiological variation in these heterogenous Alzheimer’s disease phenotypes as being along shared phenotypic continua spanning posterior cortical atrophy and Alzheimer’s disease, arising from systematic graded variations within a transdiagnostic, multidimensional neurocognitive geometry.

## Introduction

Alzheimer’s disease (AD) generates heterogeneous amnestic (typical) and non-amnestic (atypical) phenotypes ^1, 2^, including visual, logopenic, behavioural and dysexecutive presentations ^3^. Posterior cortical atrophy (PCA) includes symptoms of space and object perception deficits, constructional apraxia, environmental agnosia and alexia ^4^, and is sometimes considered a “visual-spatial AD”^5^ variant. However, considering PCA as categorically distinct from AD, i.e., adopting categorical classifications of AD variants, does not fully capture the graded variation within and between variants, or mixed phenotypes ^2, 4, 6^. This presents challenges for: diagnosing AD variants; selecting appropriate therapeutics and rehabilitation pathways; and research recruitment ^5, 6^. The current study utilised deeply phenotyped neuropsychological and neuroimaging data in AD and PCA to explore graded patient variations, rather than categorical classifications, to establish and map the neuropsychological and neuroimaging dimensions that underpin transdiagnostic (i.e., encompassing both diagnostic groups) variations in these patients.

Previous comparative studies have shown PCA and amnestic AD differ in key cognitive and visual domains (e.g., delayed auditory/verbal memory worse in amnestic AD), but not significantly in others (e.g., working memory, language, ideomotor praxis). For example, although dorsal/spatial and ventral/perceptual subtypes of PCA have been proposed ^7^, impairments in other cognitive domains are also documented, such as linguistic impairments comparable with logopenic progressive aphasia (“language-variant AD”) ^8^ and verbal short-term memory deficits found in some PCA cases, reminiscent of language-led AD ^9^. Furthermore, in amnestic (typical) AD, impairments in non-amnestic (atypical) domains including visuospatial processing have been found ^2, 10^. These findings highlight the potential for graded, overlapping cognitive variation within and between PCA and AD, which may have been missed in many studies to date that employ categorical classification systems to define groups ^9, 11^. This gap can be addressed by employing approaches that allow reconceptualising of proposed variants/subtypes of patients as occupying subregions of a graded multidimensional space, with fuzzy boundaries between ‘groups’ ^2, 10–12^ rather than discrete categorical classifications. Such approaches have been successfully applied to post-stroke aphasia^13^, primary progressive aphasia ^14^, semantic dementia ^15^, fronto-temporal lobar degeneration ^16^ and logopenic progressive aphasia ^17^. The current study therefore aimed to address this gap in AD and PCA by employing an approach which: (1) situates participants with amnestic AD and PCA in the same graded multidimensional space, rather than employing contrastive group-level statistical comparisons, to better capture the patterns of overlapping and/or non-overlapping cognitive performance, and then (2) relates the transdiagnostic phenotype dimensions to the pattern of atrophy across the whole brain, to understand how shared cognitive variation may reflect common atrophy patterns.

Using this approach, we hypothesised that we would find: (1) in AD, a dimension capturing graded variation in cognitive impairments characteristic of amnestic AD, and a dimension capturing graded variation in visuospatial impairment (as this is commonly impaired in typical AD and thus included in global dementia measures such as the MMSE and ACE-R); (2) in PCA, dimensions capturing graded variation in visuo-spatial and visuo-perceptual impairments (given the proposed dorsal/spatial and ventral/perceptual subtypes^7^), and a dimension capturing non-visual, cognitive impairments too ^9^; and (3) neural correlates for these extracted dimensions which reflect previous evidence of brain-behaviour relationships in these patient groups, e.g., occipito-parietal and occipito-temporal cortex for visuo-spatial and visuo-perceptual dimensions respectively ^18^, medial temporal lobe structures like entorhinal cortex ^19^ plus interior parietal and lateral temporal cortices for dimensions capturing diverse, non-visual impairments. Finally, given the prior evidence for overlapping phenotypic presentations within and between PCA and ‘typical’ AD, we hypothesised that there would be overlapping graded variation in PCA and AD on these extracted dimensions and that this shared cognitive variation might be reflected by common atrophy patterns in these patient groups. Specifically, our hypotheses were explored through the application of principal component analysis to a detailed neuropsychological database followed by grey matter voxel-based morphometry, allowing a data-driven exploration of (a) the presence and cognitive nature of phenotypic continua in each group; and (b) the extent of intragroup and intergroup graded variation in cognition and grey matter volume in the multidimensional space defined by these dimensions.

## Methods

### Study population

All participants were recruited at a specialist centre, the Dementia Research Centre (DRC) at the National Hospital for Neurology and Neurosurgery, London, UK. All participants in this study were first interviewed on their history of behavioural, neuropsychiatric, dementia- and non-dementia-related neurological symptoms. Participants were then identified based on the interview and documentations related to their diagnosis, such as clinical letters and summaries of their medical and symptom history. All PCA participants met consensus criteria for PCA-pure ^4^, and Tang-Wai et al. ^20^ and Mendez et al. ^21^ clinical criteria based on available information at baseline and expert retrospective clinical review. PCA participants were excluded from this study if there was evidence of non-AD dementia (i.e., DLB or CBD), including CSF/Amyloid-PET incompatible with underlying AD and/or clinical features of early visual hallucinations, pyramidal signs, reduplicative phenomena, parkinsonism, alien limb syndrome, asymmetric dystonia and myoclonus and ataxia. All AD participants met the National Institute of Neurological and Communicative Disorders and Stroke and the Alzheimer’s Disease and Related Disorders Association (NINCDS-ADRDA) criteria for probable AD with recently proposed revisions ^22^. AD participants were excluded if they showed a non-amnestic presentation consistent with the diagnostic criteria for atypical Alzheimer’s disease (posterior cortical atrophy, logopenic progressive aphasia, corticobasal syndrome, or behavioural/dysexecutive AD). Consequently, this group consisted of participants with amnestic-led AD presentations.

All available molecular or pathological evidence (34 PCA; 39 AD) supported underlying AD pathology (63 had a CSF profile compatible with AD), 3 had positive amyloid PET scans; 11 had autopsy-proven Alzheimer’s disease. Patients with biomarker evidence of Alzheimer’s disease pathology met the McKhann *et al.* ^22^ IWG-2 criteria for probable Alzheimer’s disease with high biomarker probability of Alzheimer’s disease aetiology ^23^.

The PCA and AD cases have been included in previous publications ^12, 19, 24^. All patients provided informed consent under approval from NRES Committee London, Queen Square.

### Neuropsychological assessments

Both groups completed the same neuropsychological battery, thus allowing direct comparisons. The neuropsychological assessments were completed typically on the same day as the neuroimaging scan, or where this was not possible, the scan and neuropsychological assessments took place within 3-6 months of each other. The tests included in the principal component analysis are shown in Table 2 and most are described in Lehmann *et al.* ^24^, with the addition of letter “A” Cancellation ^25^, recognition memory for faces ^26^, and tests of early visual processing. The latter included: hue discrimination (CORVIST ^27^), shape discrimination ^28^, figure/ground separation (VOSP – Visual Object and Space Perception battery ^29^), and crowding. Assessments measuring time to complete or number of errors, where a lower value indicates less impaired performance, were inverted so that lower values across all tests indicated worse performance. Significant differences between diagnostic groups on each neuropsychological test were assessed through independent t-tests.

### Cognitive analysis

All raw cognitive scores were converted to percentages. For time-based measures without a fixed maximum score (letter ‘A’ cancellation (time); Crowding (time); VOSP dot count (time)), scores were converted to a percentage of the maximum time taken within each cohort. The adequacy of the sample size for each principal component analysis was determined using the Kaiser-Meyer-Olkin measure of sampling adequacy and Bartlett’s test of sphericity.

#### Imputation and component selection

To retain as much information (patients and tests) as possible, missing data were imputed using probabilistic principal component analysis (PPCA) ^30^, which was also used to select the optimal number of components for subsequent principal component analysis using the imputed dataset (as described in Ingram *et al.* ^14^, see Supplement). The subsequent principal component analyses were also run on a version of the dataset with missing data more strictly removed (see Supplement).

#### Principal component analysis

We applied separate principal component analyses to the AD and PCA cohorts to establish the multidimensional space of each presentation independently (this avoids the possible danger of creating false overlaps by fusing the two groups into an unrepresentative single homogenous space). The principal component analysis for the AD group is shown in the Supplement. We applied varimax rotation to promote cognitive interpretation of the emergent dimensions (as well as comparisons across the two multidimensional spaces). Normalised factor scores were obtained for each patient, for subsequent neuroimaging analyses and creation of the scatterplots.

Having established the multidimensional spaces for AD and PCA independently, we then explored whether there were any regions of these multidimensional spaces showing transdiagnostic overlap in impairments. This was achieved by projecting the neuropsychological scores from one group through the coefficient matrix of the other group (as both cohorts underwent the same cognitive test battery). The results obtained by projecting PCA patients into the AD-derived multidimensional space are presented in Figure 1 Panel D (the AD principal component analysis is presented in full in the Supplement). We also explored whether the extracted components were related to disease severity (see Supplement).

**Figure 1.**
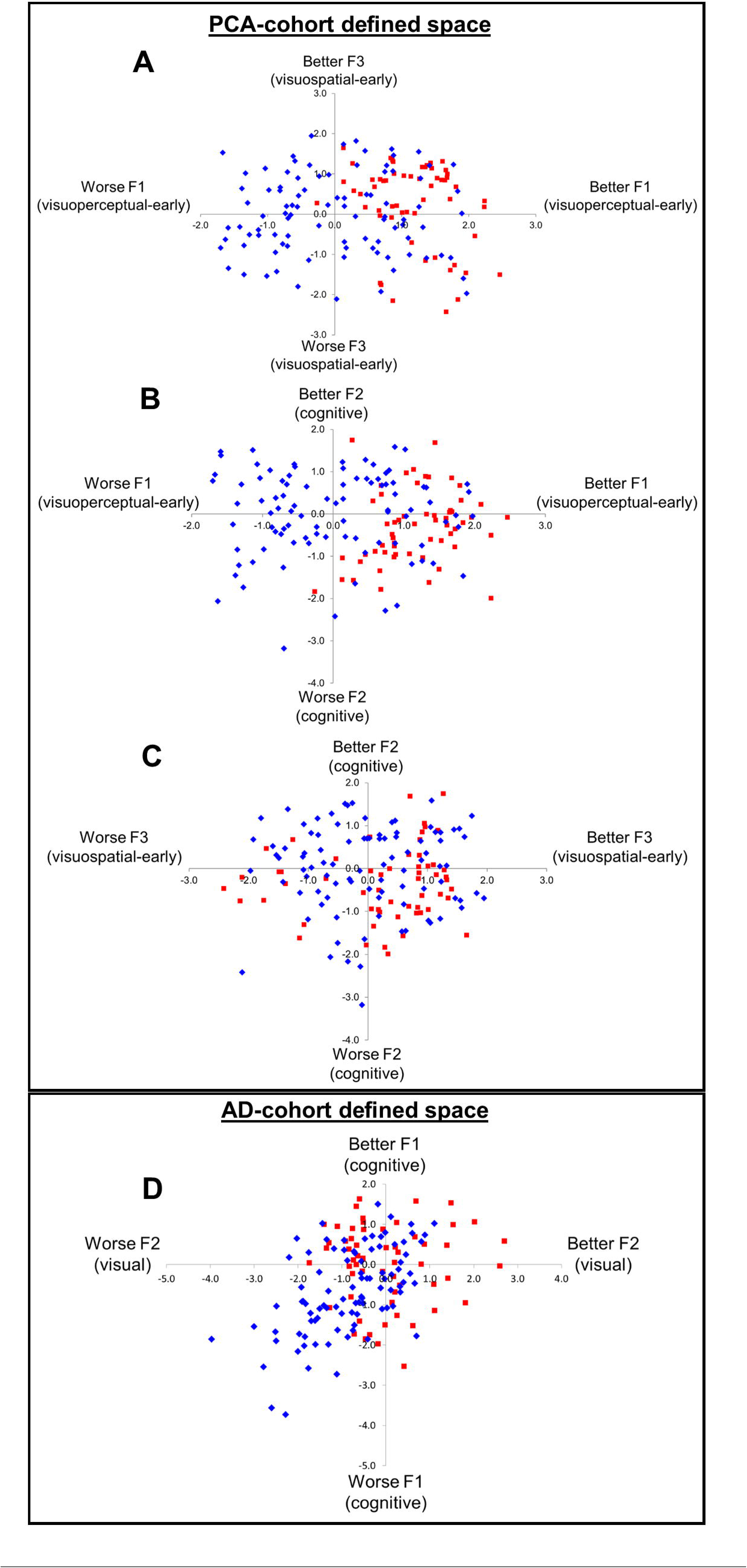
Graded intergroup phenotypic variation in posterior cortical atrophy and Alzheimer’s disease. Panels A-C: Alzheimer’s disease (AD) cases projected into posterior cortical atrophy (PCA) multidimensional space. Panel D: PCA cases projected into AD multidimensional space. Key: AD – red squares; PCA - blue diamonds.

#### Image Acquisition

T1-weighted volumetric MR scans were acquired for 71 healthy controls, 70 PCA patients and 14 AD patients over a 10-year period from 2005 – 2015. Seven PCA and 5 AD scans were excluded after image quality assurance due to motion and ghosting artifacts, yielding a total number of 71 healthy control, 62 PCA and 9 AD scans that were included in the final analyses. The majority of scans (controls: 39; PCA: 43; AD: 8) were acquired on a Siemens Prisma 3T scanner using a magnetisation prepared rapid acquisition gradient echo sequence (MPRAGE) with a 256×256 acquisition matrix, 282mm field of view and the following acquisition parameters: TE=2.9ms, TR=2200ms and TI=900ms. The remaining images (controls: 32; PCA: 19; AD: 1) were acquired on a 1.5T Sigma MRI scanner using a spoiled gradient echo (SPGR) sequence with a 256×256 image matrix, a 240mm field of view and the following acquisition parameters: TE=6.3ms; TR=14.2ms and TI=650ms.

#### Image Pre-processing

Image pre-processing involved the following steps conducted using Statistical Non-Parametric mapping (SnPM - ^31^ – a toolbox within Statistical Parametric Mapping software (SPM12.1 -)) (1) image format conversion from DICOM to NIFTI; (2) tissue segmentation using SPM’s unified model ^32^ (3) the creation of a study-specific grey matter (GM) segment template using SHOOT; (4) normalisation of the segments to the study specific template that generally matches standard space (MNI) in orientation using SHOOT transformations; (5) modulation to account for local volume changes; and (6) smoothing using a 6 mm full-width at half-maximum Gaussian kernel to compensate for inaccuracies in spatial alignment and between-subject differences in anatomy. The smoothed, normalised and modulated SHOOT-imported GM segments were then used for analysis. Image pre-processing steps (3)-(6) were performed for the different analyses (PCA-only and Combined) separately to ensure that the GM segment template only included analysis-specific participant scans.

### Voxel-based morphometry

We used whole-brain voxel-based morphometry (VBM) to explore the relationship between brain atrophy and graded variation in cognitive performance in PCA and AD. VBM analysis was performed using Statistical Non-Parametric mapping (SnPM ^31^ using SPM12.1) which allows for pseudo t-statistic images to be assessed for significance using a standard non-parametric multiple comparisons procedure based on permutation testing. Prior to performing the analyses, a whole-brain GM mask was defined to include only voxels for which the intensity was at least 0.2 in at least 80% of the images to circumvent exclusion of voxels most vulnerable to brain atrophy ^33^.

#### Correlations Between Grey Matter Volume and Principal Component-Derived Factor Scores

Two VBM regression analyses were performed using factor scores from the PCA-derived multidimensional space, a PCA-only (N=62) and a PCA/AD combined (N=71) analysis to explore PCA-specific and shared PCA/AD associations between GM volume and neuropsychological deficits, respectively. The Combined VBM analysis used factor scores from the PCA principal component analysis, either directly (for PCA cases) or through projecting raw neuropsychological scores through the PCA-derived coefficient matrix (for AD cases), to relate variation in the same multidimensional space to GM volume across both groups. Both regression models included smoothed, modulated, and warped GM volume as the dependent variable, the three PCA principal component-generated factor scores as the independent variables, and age at assessment (mean-centred), total intracranial volume (mean-centred), gender and scanner (3T or 1.5T) as covariates. The Combined VBM analysis included group as an additional covariate. An AD-only analysis (i.e., relating GM volume to factor scores from the AD-derived multidimensional space with projected scores for PCA cases) was not performed due to the limited number of available AD scans.

Statistical significance was determined by permutation testing (10,000 permutations) based on peak-voxel inference and set at p<.05 (family-wise error corrected). Scatterplots were created to visualise the relationship between GM volume and factor scores. The 3D volume results were projected to the surface using MRIcroGL (version 14 - https://www.nitrc.org/projects/mricrogl).

#### Grey Matter Volume Changes in Posterior Cortical Atrophy and Alzheimer’s Disease

To aid interpretation of correlation analyses, we assessed differences in voxel-wise GM volume in PCA and AD relative to healthy controls separately using independent t-tests. Age at assessment (mean-centred), total intracranial volume (mean-centred), gender, and scanner (3T or 1.5T) were included as covariates. Effect size maps are presented in Supplementary Materials.

### Data Availability

Anonymized data associated with this article will be made available by request from any qualified investigator.

## Results

### Patients

Ninety-three people with PCA and 58 people with AD were included in this study. Demographic details are summarised in Table 1. There were no significant differences between the AD and PCA groups in either age (t_(137)_ = .569, *p* = .571) or symptom duration (t_(115)_ = 1.907, *p* = .059). There were more females than males in the PCA group, and more males than females in the AD group (χ^2^_(1)_ = 9.35, p = .002). MMSE scores were not significantly different between AD and PCA (t_(141)_ = -1.73, *p* = .085).

**Table 1.** Demographic details for each diagnostic group. Age, symptom duration, MMSE score and Total Intracranial Volume (TIV) are presented as mean (SD). The total sample size per group is given in “Total N” with the number of females in the group given in brackets (F). The sample size for TIV is 62 PCA, 9 AD.

| Diagnosis | Total N<br>(F) | Age<br>(years) | Symptom duration<br>(years) | MMSE | Total Intracranial Volume<br>(mm<sup>3</sup>) |
| --- | --- | --- | --- | --- | --- |
| AD | 58 (22) | 57.1 (6.4) | 6.2 (3.0) | 19.7<br>(4.9) | 1422.7 (134.1) |
| PCA | 93 (59) | 59.9 (8.1) | 5.2 (2.6) | 21.2<br>(5.1) | 1439.1 (158.3) |

### Neuropsychological tests

Scores on all neuropsychological tests for AD and PCA participants are summarised in Table 2.

**Table 2.**
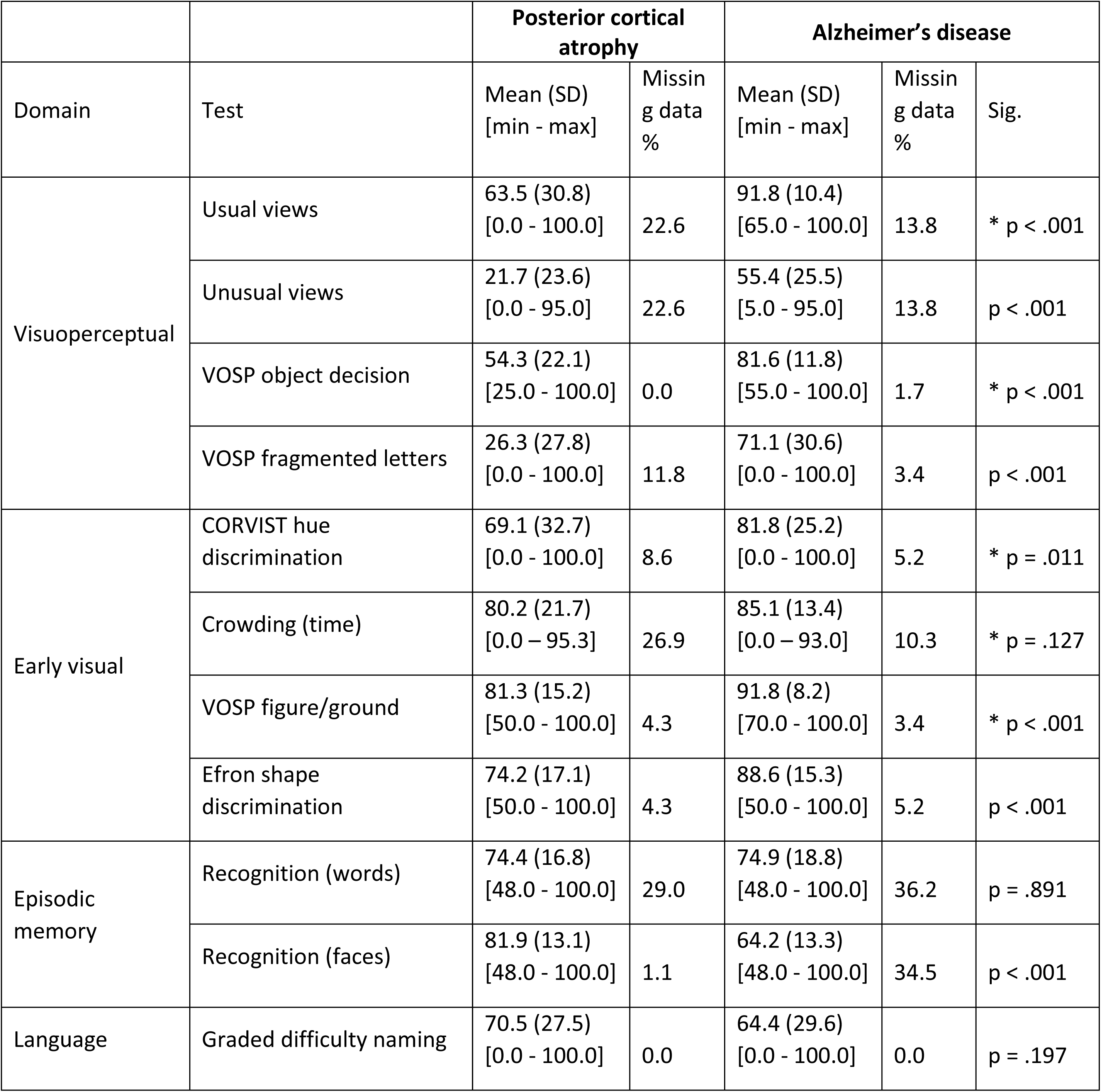
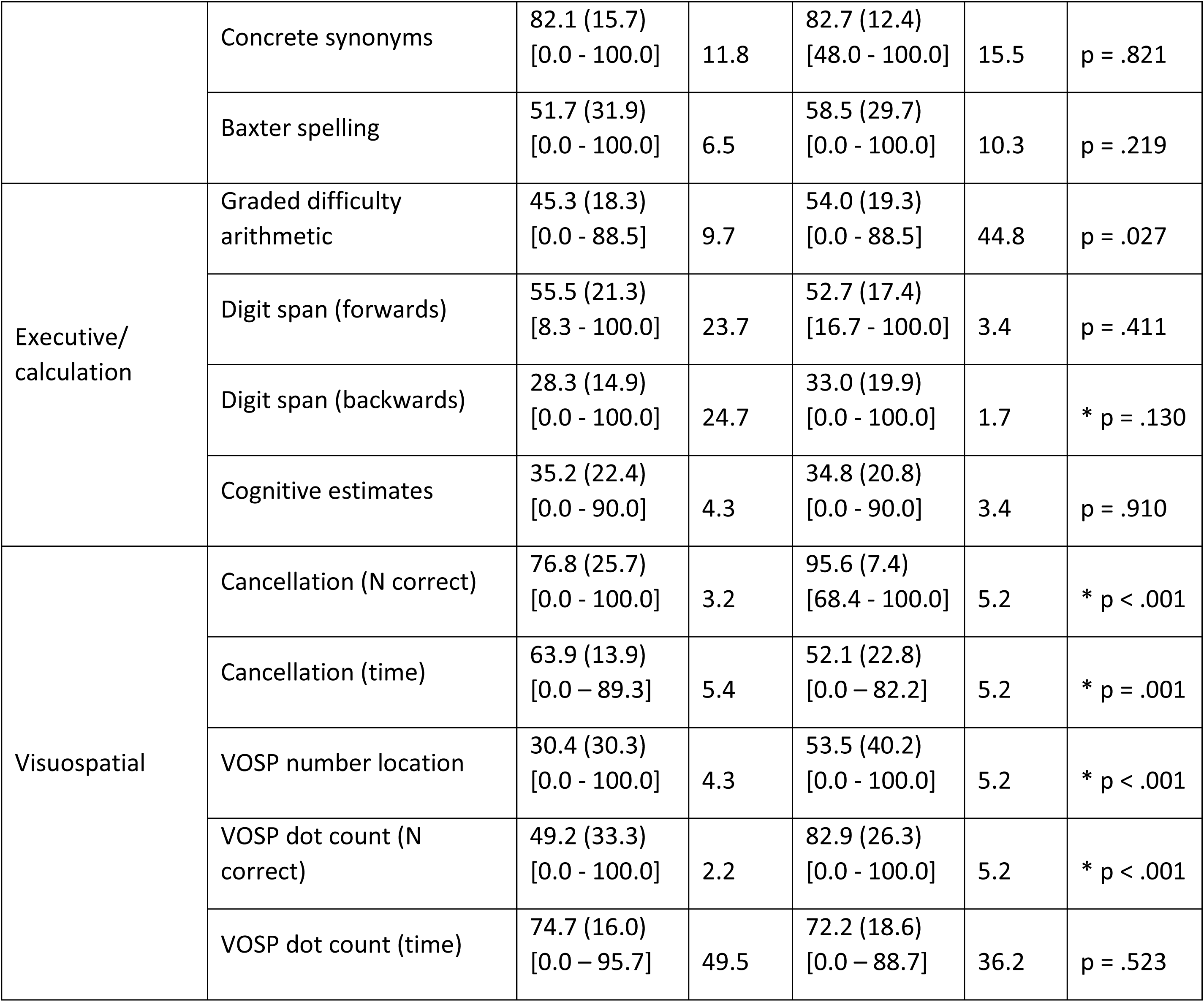
Neuropsychology test scores and missing data. Neuropsychology test scores shown as percentage of maximum score per group (higher percentage corresponds to less impairment (less errors, faster time to complete). Missing data is shown as percentage missing per group. Significant differences between diagnostic groups on each test assessed through independent t-tests; * Mann-Whitney U statistic reported due to heterogeneity of variance. Abbreviations: SD – standard deviation; VOSP – Visual Object and Space Perception; CORVIST – Cortical Vision Screening Test.; N correct – number of items correct.

| Domain | Test | Posterior cortical atrophy |  | Alzheimer's disease |  |  |
| --- | --- | --- | --- | --- | --- | --- |
|  |  | Mean (SD)<br>[min - max] | Missing data<br>% | Mean (SD)<br>[min - max] | Missing data<br>% | Sig. |
| Visuoperceptual | Usual views | 63.5 (30.8)<br>[0.0 - 100.0] | 22.6 | 91.8 (10.4)<br>[65.0 - 100.0] | 13.8 | * p < .001 |
|  | Unusual views | 21.7 (23.6)<br>[0.0 - 95.0] | 22.6 | 55.4 (25.5)<br>[5.0 - 95.0] | 13.8 | p < .001 |
|  | VOSP object decision | 54.3 (22.1)<br>[25.0 - 100.0] | 0.0 | 81.6 (11.8)<br>[55.0 - 100.0] | 1.7 | * p < .001 |
|  | VOSP fragmented letters | 26.3 (27.8)<br>[0.0 - 100.0] | 11.8 | 71.1 (30.6)<br>[0.0 - 100.0] | 3.4 | p < .001 |
| Early visual | CORVIST hue discrimination | 69.1 (32.7)<br>[0.0 - 100.0] | 8.6 | 81.8 (25.2)<br>[0.0 - 100.0] | 5.2 | * p = .011 |
|  | Crowding (time) | 80.2 (21.7)<br>[0.0 - 95.3] | 26.9 | 85.1 (13.4)<br>[0.0 - 93.0] | 10.3 | * p = .127 |
|  | VOSP figure/ground | 81.3 (15.2)<br>[50.0 - 100.0] | 4.3 | 91.8 (8.2)<br>[70.0 - 100.0] | 3.4 | * p < .001 |
|  | Efron shape discrimination | 74.2 (17.1)<br>[50.0 - 100.0] | 4.3 | 88.6 (15.3)<br>[50.0 - 100.0] | 5.2 | p < .001 |
| Episodic memory | Recognition (words) | 74.4 (16.8)<br>[48.0 - 100.0] | 29.0 | 74.9 (18.8)<br>[48.0 - 100.0] | 36.2 | p = .891 |
|  | Recognition (faces) | 81.9 (13.1)<br>[48.0 - 100.0] | 1.1 | 64.2 (13.3)<br>[48.0 - 100.0] | 34.5 | p < .001 |
| Language | Graded difficulty naming | 70.5 (27.5)<br>[0.0 - 100.0] | 0.0 | 64.4 (29.6)<br>[0.0 - 100.0] | 0.0 | p = .197 |
|  | Concrete synonyms | 82.1 (15.7)<br>[0.0 - 100.0] | 11.8 | 82.7 (12.4)<br>[48.0 - 100.0] | 15.5 | p = .821 |
|  | Baxter spelling | 51.7 (31.9)<br>[0.0 - 100.0] | 6.5 | 58.5 (29.7)<br>[0.0 - 100.0] | 10.3 | p = .219 |
| Executive/<br>calculation | Graded difficulty<br>arithmetic | 45.3 (18.3)<br>[0.0 - 88.5] | 9.7 | 54.0 (19.3)<br>[0.0 - 88.5] | 44.8 | p = .027 |
|  | Digit span (forwards) | 55.5 (21.3)<br>[8.3 - 100.0] | 23.7 | 52.7 (17.4)<br>[16.7 - 100.0] | 3.4 | p = .411 |
|  | Digit span (backwards) | 28.3 (14.9)<br>[0.0 - 100.0] | 24.7 | 33.0 (19.9)<br>[0.0 - 100.0] | 1.7 | * p = .130 |
|  | Cognitive estimates | 35.2 (22.4)<br>[0.0 - 90.0] | 4.3 | 34.8 (20.8)<br>[0.0 - 90.0] | 3.4 | p = .910 |
| Visuospatial | Cancellation (N correct) | 76.8 (25.7)<br>[0.0 - 100.0] | 3.2 | 95.6 (7.4)<br>[68.4 - 100.0] | 5.2 | * p < .001 |
|  | Cancellation (time) | 63.9 (13.9)<br>[0.0 - 89.3] | 5.4 | 52.1 (22.8)<br>[0.0 - 82.2] | 5.2 | * p = .001 |
|  | VOSP number location | 30.4 (30.3)<br>[0.0 - 100.0] | 4.3 | 53.5 (40.2)<br>[0.0 - 100.0] | 5.2 | * p < .001 |
|  | VOSP dot count (N<br>correct) | 49.2 (33.3)<br>[0.0 - 100.0] | 2.2 | 82.9 (26.3)<br>[0.0 - 100.0] | 5.2 | * p < .001 |
|  | VOSP dot count (time) | 74.7 (16.0)<br>[0.0 - 95.7] | 49.5 | 72.2 (18.6)<br>[0.0 - 88.7] | 36.2 | p = .523 |

### Establishing the multidimensional spaces of Posterior Cortical Atrophy and Alzheimer’s Disease

The principal component analysis for the PCA group was robust (Kaiser-Meyer-Olkin = 0.865) and Bartlett’s test of sphericity was significant (approximate χ^2^ = 1242.972, d.f. = 231, *p* < 0.001). The 3-factor varimax rotated solution accounted for 61.0% of the total variation in the patients’ performance. The variance explained per factor is as follows: Factor 1 (visuoperceptual-early) = 23.0%; Factor 2 (cognitive) = 21.4%; Factor 3 (visuospatial-early) = 16.6%. The factor loadings are shown in Table 3. A summary of tests loading onto each factor and hence the term used to label each factor is presented in the Supplement, with tests for the relationship of each factor with disease severity. This multidimensional space was used for the following analyses, so the principal component analysis result for the AD group alone is shown in eTable 1 in the Supplement.

**Table 3.** Principal component analysis results for posterior cortical atrophy. Factor loadings larger than 0.5 are shown in bold. Abbreviations: VOSP – Visual Object and Space Perception; CORVIST – Cortical Vision Screening Test; N correct – number of items correct.

| Domain | Test | Factor 1<br>(visuoperceptual-early) | Factor 2<br>(cognitive) | Factor 3<br>(visuospatial-early) |
| --- | --- | --- | --- | --- |
| Visuo-perceptual | Usual views | **0.894** | 0.070 | 0.244 |
|  | Unusual views | **0.871** | -0.037 | -0.021 |
|  | VOSP object decision | **0.857** | 0.017 | 0.135 |
|  | VOSP fragmented letters | **0.648** | 0.115 | 0.464 |
| Early visual | CORVIST hue discrimination | **0.627** | 0.198 | 0.246 |
|  | Crowding (time) | **0.655** | 0.222 | 0.429 |
|  | VOSP figure/ground | **0.526** | 0.021 | 0.460 |
|  | Efron shape discrimination | 0.454 | 0.125 | 0.413 |
| Episodic memory | Recognition (words) | **0.770** | 0.131 | 0.206 |
|  | Recognition (faces) | -0.098 | **0.548** | 0.258 |
| Language | Graded Difficulty Naming | 0.193 | **0.781** | -0.036 |
|  | Concrete synonyms | 0.135 | **0.773** | 0.103 |
|  | Baxter spelling | 0.113 | **0.795** | 0.165 |
| Executive/calculation | Graded Difficulty Arithmetic | 0.011 | **0.743** | 0.342 |
|  | Digit span (forwards) | 0.128 | **0.702** | 0.077 |
|  | Digit span (backwards) | -0.052 | **0.803** | 0.042 |
|  | Cognitive estimates | -0.202 | **-0.684** | -0.225 |
| Visuo-spatial | Cancellation (N correct) | 0.365 | 0.305 | **0.589** |
|  | Cancellation (time) | 0.252 | 0.220 | **0.542** |
|  | VOSP number location | 0.237 | 0.147 | **0.814** |
|  | VOSP dot count (N correct) | 0.163 | 0.037 | **0.823** |
|  | VOSP dot count (time) | 0.186 | 0.318 | **0.648** |

### Phenotypic continua in Posterior Cortical Atrophy and Alzheimer’s disease

We explored whether PCA and AD cases overlapped with each other in their respective multidimensional spaces, by projecting factor scores of one group into the multidimensional space of the other. AD cases (red squares) projected into the PCA-defined space are shown in Figure 1A-C, whilst PCA cases (blue diamonds) projected into AD-defined space are shown in Figure1D.

These comparative plots illustrate some key observations: (i) there are graded variations along all dimensions in both patient groups; (ii) there is considerable overlap between the AD and PCA groups on the general cognitive impairment dimension, irrespective of which principal component analysis solution is used; (iii) the AD also overlap with the PCA group in terms of the visuospatial and visuoperceptual dimensions extracted by the PCA-cohort analysis (upper-right quadrants of Figure1A; and the right halves of B & C) – again pointing to the observation that the symptomatology of the two groups overlap; (iv) whilst a subset of the PCA cases overlap with the AD cases, there are PCA cases with more pronounced visuospatial and/or visuoperceptual impairment than AD at the same level of generalised cognitive impairment.

### Shared neural correlates of cognition across phenotypes

Regional reductions in GM volume in PCA and AD relative to control groups were consistent with previous investigations (Supplemental eFigure 2). A detailed summary of the PCA VBM results can be found in eTable 2 in the Supplement. To explore the overlapping visual and cognitive profiles in the PCA-cohort multidimensional space, these profiles were related to underlying neuroanatomy in the Combined VBM. Figure 2 shows the results of this combined analysis including PCA and AD cases with available scans.

**Figure 2.**
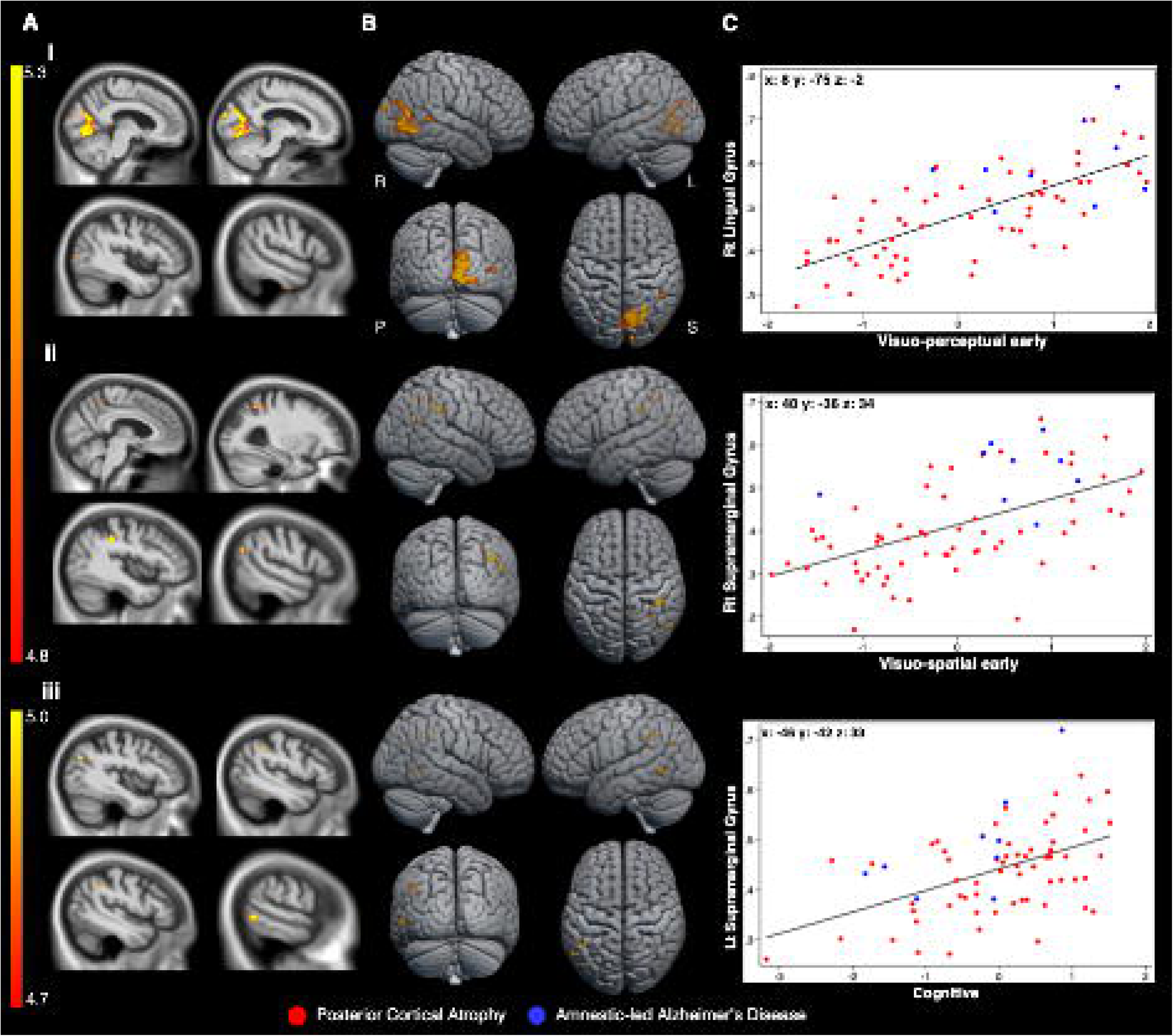
Whole-Brain VBM Results in PCA & AD. Presented are significant positive associations between neuropsychological performance and GM volumes in PCA and AD. FWE-corrected significant p < .05 regions, identified by permutation-based peak-voxel inference, are shown A) overlaid on 2-dimensional orthogonal sagittal slices of the normalised study-specific T1-weighted group average, B) surface rendered; and C) shows correlations between neuropsychological scores and participant-specific mean cluster GM volume values (largest significant cluster) by group as scatterplots. Colour bar represents t-values. MNI coordinates (mm) at peak voxel are shown in bold. Rt: right; Lt: left; i: visuoperceptual-early factor, ii: visuospatial-early factor and iii: cognitive factor; R: right; L: left; S: superior; P: posterior.

In line with the combined analysis comprising mostly PCA participant scans (PCA n=62; AD n=9), associations between factors and regional GM volume are broadly consistent with analyses restricted to the PCA group (see Supplement). To visualise the relationship between shared neural correlates of the overlapping neuropsychological variation, Figure 2Bi-iii shows, for the largest cluster associated with each principal component, the GM volume in the cluster against the corresponding factor score for every patient. This shows graded variation with and between the AD and PCA cases, for example with several AD participants exhibiting scores on visuoperceptual-early factors and lingual gyral atrophy which are commensurate with PCA group mean scores/atrophy. Additional correlates identified through combined analysis include lower visuospatial-early factor scores being associated with precuneal GM decreases (Table 4). These may relate to neuropsychological deficits and atrophy patterns (for example, diminished visuospatial functioning and precuneal atrophy) which are common across PCA and AD, particularly given the relatively young age of our AD sample. Overall, these results show graded, transdiagnostic phenotypic dimensions that relate to common atrophy patterns in these presentations of AD.

**Table 4.** Combined VBM results showing posterior cortical atrophy and Alzheimer’s disease shared brain regions in which GM volume reductions were associated with lower visuo-perceptual, visuo-spatial and cognitive factor scores. Rt: right; Lt: left; k: cluster size; PFWE: Family-wise error corrected p-value p<.05 ; x, y, z: peak-voxel MNI coordinates.

|  | k | P<sub>FWE</sub> | T | x | y | z | Brain Region |
| --- | --- | --- | --- | --- | --- | --- | --- |
| Visuoperceptual-early | 2038 | .0003 | 6.66 | 8 | -75 | -2 | Rt Lingual Gyrus |
|  |  | .0003 | 6.64 | 10 | -74 | 20 | Rt Intracalcarine Cortex |
|  |  | .0003 | 6.48 | 11 | -69 | -4 | Rt Occipital Fusiform Gyrus |
|  | 27 | .0014 | 5.99 | 42 | -93 | 20 | Rt Occipital Pole |
|  | 26 | .0050 | 5.64 | 51 | -80 | 8 | Rt Lateral Occipital Cortex |
|  | 58 | .0169 | 5.26 | 36 | -46 | 6 | Rt Medial Temporal Gyrus |
| Visuospatial-early | 120 | .0035 | 5.78 | 40 | -36 | 34 | Rt Supramarginal Gyrus |
|  | 44 | .0020 | 6.02 | 50 | -62 | 24 | Rt Lateral Occipital Cortex |
|  | 28 | .0140 | 5.28 | 30 | -54 | 44 | Rt Superior parietal lobule |
|  | 21 | .0186 | 5.18 | 46 | -28 | 30 | Rt Parietal Operculum |
|  | 17 | .0139 | 5.28 | 6 | -46 | 50 | Rt Precuneus |
| Cognitive | 69 | .0051 | 5.56 | -46 | -42 | 33 | Lt Supramarginal Gyrus |
|  | 60 | .0067 | 5.46 | -42 | -48 | 38 | Lt Angular Gyrus |
|  | 60 | .0107 | 5.30 | -56 | -57 | -2 | Lt Medial Temporal Gyrus |
|  | 55 | .0411 | 4.85 | -45 | -46 | 18 | Lt Inferior Temporal Gyrus |
|  | 24 | .0143 | 5.20 | -39 | -70 | -28 | Lt Lateral Occipital Cortex |

## Discussion

The presence of AD phenotypic variations poses particular challenges for correct diagnosis and clinical management ^5, 6^. This data-driven comparison of PCA and AD allowed us to consider to what extent varying presentations of AD are separable, mutually exclusive clinical categories or gradedly-different positions within a single, transdiagnostic (i.e., encompassing both diagnostic groups) multidimensional space. We subsequently explored whether the cognitive impairments demonstrated in PCA and AD were associated with the same neural correlates (or could be driven by atrophy in disparate brain regions). The current study provides evidence of overlapping features (visual, cognitive, and posterior cortical) in a deeply phenotyped sample of PCA and AD participants administered the same detailed neuropsychological battery. These novel comparisons extend work investigating variation within PCA ^34^ and AD ^35^, separately.

The results were broadly consistent with the conceptualisation of AD and PCA as varying continuously on a spectrum of cognitive-neuroanatomical changes: (1) both AD and PCA data generated dimensions of graded and not clustered variation in terms of generalised cognitive and visual impairments; (2) there was considerable overlap of the two patient groups along these dimensions, (3) the relationship between cognitive impairments and underlying regions of brain atrophy in PCA persisted in AD. In the remainder of the Discussion, we will consider the graded nature of the identified phenotypic variations and the implications for future clinical research and practice.

### Continua of visual processing impairment and cognitive status

Plotting PCA and AD in the respective multidimensional space from the principal component analysis demonstrated graded variation within and between these groups with respect to visual processing impairments. As expected, a good proportion of the PCA patients had more severe visuospatial and/or visuoperceptual impairments than the AD cases. However, there was a subset of AD cases who overlapped with PCA cases on the visual processing dimensions (Figure1C), indicating visual deficits commensurate with mild to moderate PCA. This finding aligns with previous early reports of AD cases with pronounced visual processing deficits ^2^ and recent findings suggesting a substantial proportion of ‘typical’ AD patients exhibit predominant visuospatial deficits ^36^. Although visual processing impairments are not necessary or sufficient for diagnosis of ‘typical’ AD, it is generally recognised that visuospatial deficits can be present or emerge later ^37^. In our sample of amnestic-led AD cases, the profile of AD cases with visual deficits commensurate with mild to moderate PCA was not confined to AD cases with globally poor performance; some AD cases presented with impaired visual processing even when their general cognitive status was better than most other cases (top left quadrant of Figure1C). Overall, these findings provide support for the core hypothesis for this PCA and AD comparison study, namely that both within and between presentations of AD and PCA, there is evidence of graded variation along phenotypic continua. Specifically, there is evidence of a graded dimension of visual impairment that is independent of variation in general cognitive status.

In addition to the overlap in visual processing impairments, considerable overlap of AD and PCA on the emergent ‘cognitive’ dimensions reiterates the importance of non-visual impairments in PCA ^4^. Others have found language deficits in early to intermediate stage PCA ^38^ consistent with logopenic progressive aphasia, and there is increasing evidence of both executive deficits in PCA ^39^ and frontal tau accumulation in PCA over time ^40^. The shared variations in linguistic or executive domains captured by the PCA, align with a transdiagnostic re-consideration of AD and its atypical subtypes ^2, 10, 17^ as reflecting graded involvement of different cognitive domains, rather than discrete subtypes with isolated impairments in select domains. These results also highlight the importance of fully characterising cognitive impairments in PCA because non-visual symptoms could contribute to the misdiagnosis of PCA^5, 7^.

### Shared neurodegenerative origins of cognitive impairments

The results of the combined VBM analysis suggest that atrophy in the extracted clusters is associated with impairment along the extracted cognitive dimensions, regardless of diagnostic group. Neuroimaging findings imply that overlapping cognitive features in these forms of dementia may arise from atrophy in similar brain regions. This supports the conceptualisation of PCA and AD as being within a shared, multidimensional phenotypic space, perhaps relating to graded neurodegeneration of functional brain networks, rather than as discrete subtypes caused by Alzheimer’s disease pathology (for a parallel proposal for the overlapping variations of logopenic progressive aphasia and AD, see: Ramanan *et al.* ^17^).

### Implications of graded variation

Our results indicate that a simple categorical distinction between AD and PCA based on diagnostic criteria would fail to capture the evident graded differences between these phenotypes. This raises the issue of how to relate graded, multidimensional approaches to traditional, categorical classification systems ^13^. The latter provide a useful diagnostic short-hand for clinicians and may be useful for contrastive group-level analysis. We are not proposing that the diagnostic labels should be abandoned entirely. Rather, being able to place cases from different diagnostic categories into a shared, transdiagnostic multidimensional space can highlight key intra- and inter-subgroup variations, enhancing our understanding of the diagnostic categories themselves. This approach is able to capture both graded phenotypic variation, including more atypical examples and mixed cases, as well as highlight more category-like phenotypes if they are present ^14^. Thus, a comprehensive ‘picture’ of an individual patient could include their broad label and their nuanced multidimensional profile. From a research perspective, this multidimensional approach allows for (and in fact necessitates) a more inclusive recruitment strategy which captures not only the “pure” prototypical cases but the majority of patients, who show graded phenotypic variation.

Possible clinical ramifications include identification of: (1) transdiagnostic, potentially treatable symptoms that would otherwise not be evident from research which studies only prototypical cases; and (2) graded clinico-radiological dimensions also open up the possibility of new approaches to stratification of cases for treatments, dosage titration, and other elements of clinical trials research that are based on scalar rather than categorical variations.

### Limitations

Three methodological considerations are important to acknowledge: availability of molecular/pathological evidence, scanner variation and sample sizes for VBM, and age of AD participants. Although the cohorts in this study met the respective neuropsychological criteria for AD and PCA, molecular/pathological evidence of AD was only available for a subset of cases. While all available molecular or pathological evidence (34 PCA; 39 AD) supported underlying Alzheimer’s disease pathology and patients overall were relatively young (AD: 57yrs +/- 6; PCA 60yrs +/- 8), we acknowledge that we cannot rule out contributions of non-AD pathology. We do note however that for all PCA patients who have made it to autopsy (N=11), all had a primary neuropathological diagnosis of AD.

In terms of the VBM analysis, the imaging data were acquired on scanners of different magnetic strengths, so there is a risk that our findings could be influenced by scanner-specific factors. However, covariates for scanner were regressed out after the estimation of regional brain volumes, to separate out scanner-specific biases (over/under estimation of GM due to scanner), reducing this risk. Additionally, the significantly larger proportion of scans from PCA cases (62 vs. 9 scans) for combined VBM analysis could have meant that these results were driven by associations in the PCA group, which may limit generalisability.

The AD participants were relatively young, as noted above. Younger onset AD (YOAD) patients can be more likely to have a predominant non-memory impairment ^11^, which could then increase the overlap with PCA or other atypical presentations in non-memory domains. Furthermore, YOAD has been found to have more precuneal atrophy and less pronounced medial temporal lobe atrophy compared to late onset AD (LOAD), even in patients who show a predominant amnestic phenotype ^41^, thus YOAD cases could potentially have a parietally-weighted neuroimaging profile that is more similar to PCA than LOAD. However, we also note that phenotypic heterogeneity is increasingly recognised in late onset AD too e.g.,^36^.

Taking these methodological considerations into account, we acknowledge that the results of this study represent an exploration of the shared variance in AD and PCA, as a test-case for exploring the multidimensional space shared by all AD phenotypes. In future work, it will be important to confirm molecular/pathological AD status in these cases to extend these findings towards understanding the heterogeneity caused by AD pathology specifically ^1^, to replicate these findings in larger samples (especially for VBM comparisons), and finally to replicate these findings in samples of individuals meeting criteria for LOAD to explore the potential impact of age at onset on the shared variation.

### Future directions

The current study shows that this test-case exploration of phenotypic continua in AD and PCA has promise for uncovering the nature of variation between different clinical presentations of AD. Future research could extend this beyond amnestic-led AD and PCA, to explore (i) the full extent of variability in all clinical phenotypes associated with AD pathology, and (ii) variation within PCA due to different aetiologies (e.g., AD, Lewy body disease, corticobasal degeneration ^4^). Establishing the underpinning multidimensional space in these samples would then provide an alternative framework in which variations along each dimension (rather than differences between groups) can be related to the underpinning neuroimaging and neurobiological features ^13^. Building from situating amnestic-led AD and PCA within the same multidimensional symptom-atrophy space, future research could build on important earlier work ^2^, which captured graded differences between subgroups of neurodegenerative disease instead of comparing groups of cases based on their diagnostic label.

## Acknowledgements

We thank all of the patients, their families, and carers for their continued support of our research programmes. This research was supported by The Rosetrees Trust (A1699), the ERC (GAP: 670428 - BRAIN2MIND_NEUROCOMP), MRC Career Development Award to ADH (MR/V031481/1) and MRC intramural funding (MC_UU_00005/18). The Dementia Research Centre is an Alzheimer’s Research UK Co-ordinating Centre and is supported by Alzheimer’s Research UK, Brain Research Trust, and The Wolfson Foundation. This work was also supported by the NIHR Queen Square Dementia Biomedical Research Unit and by an Alzheimer’s Research UK Senior Research Fellowship [ART-SRF2010-3] and ESRC/NIHR [ES/L001810/1] and EPSRC [EP/M006093/1] grants to SC. KY is an Etherington PCA Senior Research Fellow and is funded by the Alzheimer’s Society, grant number 453 (AS-JF-18-003). In memory of Mrs Pam Southerden.

## Author contributions

RUI, GP, AH, MALR contributed to the conception and design of the study; KXXY, SC contributed to the acquisition of the data; RUI, DO, GP, AH, DC, MALR, KXXY contributed to the analysis of the data; RUI, DO, GP, AH, DC, MALR, KXXY, SC contributed to drafting the text; RUI, DO, DC, MALR, KXXY contributed to preparing the figures.

## Open access

For the purpose of Open Access, the UKRI-funded authors have applied a CC BY public copyright licence to any Author Accepted Manuscript version arising from this submission.

