## Supplement for "Graded multidimensional clinical and radiological variation in patients with Alzheimer’s disease and posterior cortical atrophy"

*Joint first authors

†Joint senior authors

### eMethods

#### *Imputation and component selection*

To retain as much information (patients and tests) as possible, missing data were imputed using probabilistic principal component analysis (PPCA) [36], which was also used to select the optimal number of components for subsequent principal component analysis using the imputed dataset (as described in Ingram *et al.* [20]). In this procedure, the number of components for PPCA to extract must be pre-specified. A k-fold cross validation approach was used to choose the component-solution (i.e., number of components) with the lowest root mean squared error (RMSE) for held-out cases over 1000 permutations [37]. The subsequent principal component analyses were also run on a version of the dataset with missing data more strictly removed (no test having more than 12% missing data and no participant having more than 18% missing data) and the factors in the resultant solutions were correlated all above 0.94 with the corresponding components from the solutions reported below.

#### *Principal Component Analysis*

We calculated Pearson correlation coefficients between scores on all extracted factors and (a) age, (b) symptom duration, and (c) MMSE scores to explore the relationship with disease severity (correcting for multiple comparisons, Bonferroni corrected criterion for significance $p < .016$).

### Results

#### *Posterior cortical atrophy principal component analysis*

Tests of visuoperceptual function (usual/unusual views, object decision and fragmented letters), early visual processing (crowding, hue discrimination, shape discrimination and figure/ground discrimination), and recognition memory for words, loaded heavily onto Factor 1. A common feature of these tests is the identification of stimuli based on their perceptual features; hence we called this factor 'visuoperceptual-early'. This component was not significantly correlated (after correction for multiple comparisons) with MMSE ($r$ = .094, $p$ = .377), symptom duration ($r$ = -.227, $p$ = .039) or age ($r$ = -0.201, $p$ = .057).

Tests of recognition memory for faces, language abilities (synonym judgement, naming, spelling), and attention/executive tasks (digit span forwards and backwards, arithmetic, and cognitive estimates), loaded heavily onto Factor 2. This component was significantly correlated with MMSE ($r$ = 0.603, p < .001) and age ($r$ = .334, $p$ = .001), but not symptom duration ($r$ = -.021, $p$ = .850). Given the nature of the test loadings and the correlation with MMSE, we termed this factor 'cognitive'.

Tests of visuospatial processing (dot counting and number location) and letter cancellation had high loadings onto Factor 3. Moderate loadings were also found for tests of early visual processing and identifying fragmented letters. This component had a significant correlation with MMSE ($r$ = 0.398, p < .001) but not age ($r$ = .039, $p$ = .712) or symptom duration ($r$ = -.079, $p$ = .479). These tests overall require scanning of the visual field/visual search, and abstraction of visuospatial relationships between stimuli. Furthermore, variance associated with early visuoperceptual aspects of some of these visual tasks was already accounted for by the first factor. Accordingly, we called Factor 3 'visuospatial-early'.

To visualise variation along these factors, all PCA-patient factor scores were plotted (blue diamonds in main Figure1A-C). These scatterplots demonstrate various important findings: (i) for all three dimensions, the patients varied in graded ways – i.e., there was no evidence of clusters implicating mutually-exclusive subtypes of PCA; (ii) the analyses added evidence to previous suggestions that there are both visuospatial and visuoperceptual presentations of PCA (i.e., some patients presented with greater impairment in one aspect of visual processing than the other, as well as many cases with dual visual deficits); (iii) there are concurrent, varying levels of generalised cognitive impairment in PCA (Figure1B&C), consistent with it being commonly associated with underlying AD.

#### *Alzheimer's disease principal component analysis*

The principal component analysis for the AD group was also robust (Kaiser-Meyer-Olkin = 0.783) and Bartlett's test of sphericity was significant (approximate $\chi^2$ = 670.952, d.f. = 231, $p$ < .001). This produced a 2-factor rotated solution which accounted for 44.3% of variance in patients' performance (Factor 1 (cognitive) = 23.9%, Factor 2 (visual) = 20.4%). The factor loadings are shown in Supplemental eTable 1.

**Supplemental eTable 1 - Principal components analysis results for Alzheimer's disease. Factor loadings larger than 0.5 are shaded in grey. Abbreviations: VOSP – Visual Object and Space Perception; CORVIST – Cortical Vision Screening Test.; N correct – number of items correct.**

| Domain | Test | Factor 1 (cognitive) | Factor 2 (visual) |
|---|---|---|---|
| Visuo-perceptual | Usual views | 0.747 | 0.076 |

| | | | |
|---|---|---|---|
| | Unusual views | 0.766 | 0.183 |
| | VOSP object decision | 0.102 | 0.381 |
| | VOSP fragmented letters | 0.440 | 0.682 |
| Early visual | CORVIST hue discrimination | 0.508 | 0.093 |
| | Crowding (time) | 0.007 | 0.436 |
| | VOSP figure/ground | 0.254 | 0.371 |
| | Efron shape discrimination | 0.183 | 0.505 |
| Episodic memory | Recognition (words) | 0.733 | -0.179 |
| | Recognition (faces) | 0.441 | 0.198 |
| Language | Graded Difficulty Naming | 0.872 | -0.047 |
| | Concrete synonyms | 0.674 | 0.343 |
| | Baxter spelling | 0.437 | 0.331 |
| Executive/calculation | Graded Difficulty Arithmetic | 0.557 | 0.441 |
| | Digit span (forwards) | -0.011 | 0.451 |
| | Digit span (backwards) | 0.357 | 0.641 |
| | Cognitive estimates | -0.699 | -0.294 |
| Visuo-spatial | Cancellation (N correct) | 0.182 | 0.395 |
| | Cancellation (time) | 0.154 | 0.756 |
| | VOSP number location | 0.571 | 0.569 |
| | VOSP dot count (N correct) | -0.007 | 0.775 |
| | VOSP dot count (time) | 0.059 | 0.898 |

Measures that loaded heavily onto the first factor were tests of recognition memory (words and faces), visuoperceptual ability (usual and unusual views), working memory (arithmetic, digit span backwards, cognitive estimates), language (naming, concrete synonyms, spelling), hue discrimination and visuospatial processing (number location). This factor was correlated with MMSE score ($r$ = .579, $p$ < .001), but not age ($r$ = .034, $p$ = .860) or symptom duration ($r$ = .108, $p$ = .543). The diverse nature of the tests which loaded onto this factor, led us to call this factor 'cognitive'. Indeed, there was overlap between the loadings on this dimension and those found for Factor 2 (cognitive) in the PCA group data. The second AD factor was associated with tests of visuospatial ability (number location, dot counting), visuoperceptual processing (fragmented letters), early visual processing (figure/ground separation, shape discrimination, crowding), and attention/executive function (digit span, letter cancellation). We called this factor 'visual' given that it was related to most of the visual tests. This factor was correlated with MMSE score ($r$ = .412, $p$ = .002) and age ($r$ = .455, $p$ = .005) but not symptom duration ($r$ = -.257, $p$ = .143).

Supplemental eFigure 1 plots the factor scores for the AD cases (red squares) (also shown in Figure 1 Panel D in the main paper). This scatterplot indicates that for AD cases: (i) there were two clear dimensions of variation; (ii) again there were only graded variations in performance and no evidence of clusters; and (iii) there were cases with poor cognitive or visual processes, or both. The presence of AD cases with more prominent visual impairment yet less pronounced general cognitive decline reinforces previous observations of patients with prominent visual/posterior features in general AD cohort evaluations, especially with younger age at onset (see Introduction).

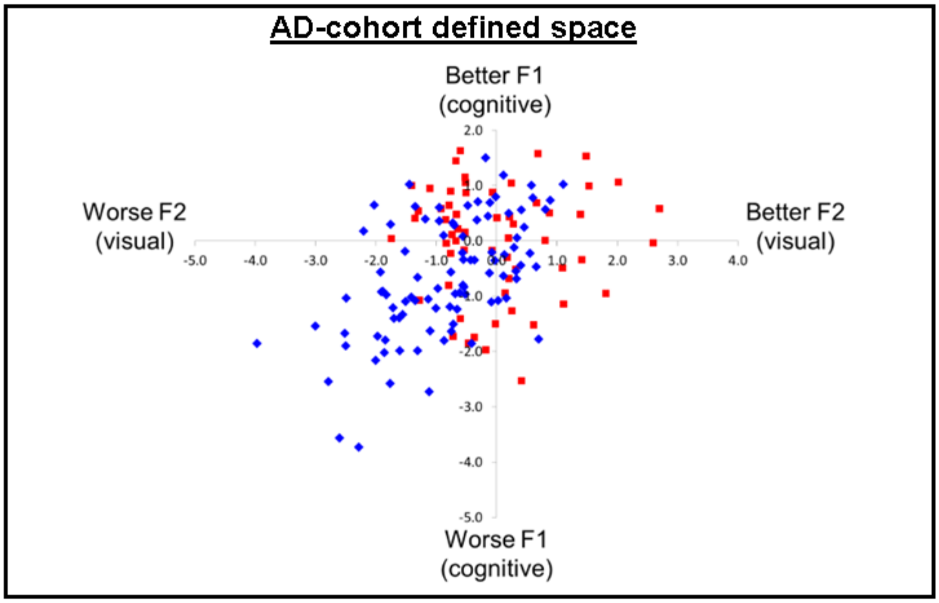

**Supplemental eFigure 1 - Graded intergroup phenotypic variation in Alzheimer's disease and posterior cortical atrophy. Posterior Cortical Atrophy (PCA) cases projected into Alzheimer's disease (AD) multidimensional space. Key: AD - red squares; PCA - blue diamonds.**

*Regional reductions in GM volume in PCA and AD relative to control groups*

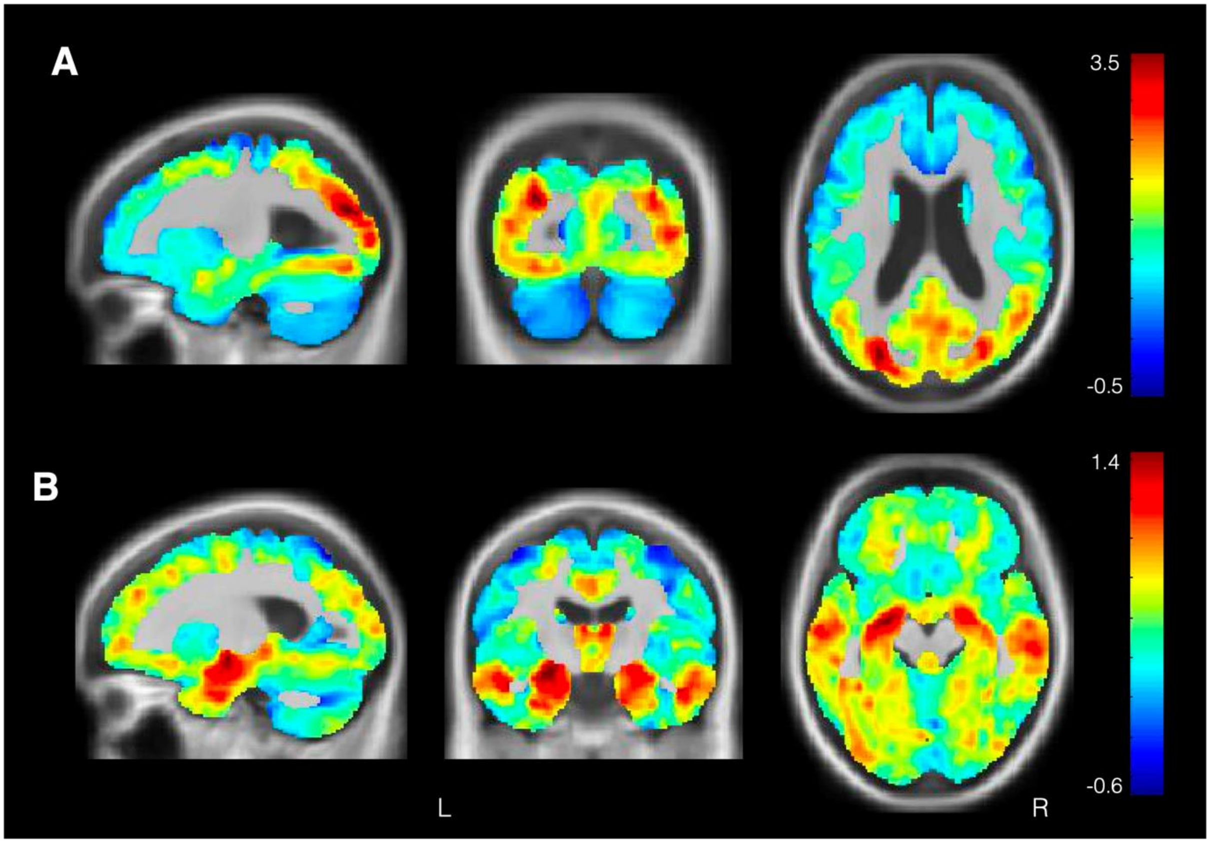

**Supplemental eFigure 2 - Whole-Brain VBM Results Grey Matter Volume Differences in PCA & AD Relative to Controls. Shown are unthresholded effect size (Cohen's d) maps overlaid on normalised study-specific T1-weighted group templates for the A) PCA < healthy controls contrast and B) AD < healthy controls contrast. Colour bar represents Cohen's d values. R: right; L: left.**

Relative to healthy controls, PCA patients displayed reductions in GM volume involving predominantly parieto-occipital regions (Supplemental eFigure 2). Other smaller clusters included the right paracingulate gyrus. Relative to healthy controls, the AD group demonstrated reduced GM volume involving predominantly temporal regions such as the bilateral hippocampus, amygdala, inferior and middle temporal gyri and fusiform cortex.

*Neural correlates of posterior cortical atrophy multidimensional space*

A detailed summary of the PCA VBM results can be found in Supplemental eTable 2.

**Supplemental eTable 2 - VBM results showing brain regions in which GM volume reductions were associated with lower visuoperceptual-early, visuospatial-early and cognitive factor scores in posterior cortical atrophy. Rt: right; Lt: left; k: cluster size; PFWE: Family-wise error corrected p-value p<.05 ; x, y ,z: peak-voxel MNI coordinates.**

| | k | $P_{FWE}$ | T | x | y | z | Brain Region |
|---|---|---|---|---|---|---|---|
| Visuoperceptual-early | 2096 | .0001 | 7.15 | 14 | -75 | 21 | Rt Cuneal Cortex |
| | | .0003 | 6.58 | 21 | -69 | -4 | Rt Occipital Fusiform Gyrus |

| | | | | | | | |
|---|---|---|---|---|---|---|---|
| | | .0006 | 6.38 | 9 | -76 | 0 | Rt Lingual Gyrus |
| | 109 | .0006 | 6.40 | -12 | -42 | -39 | Rt Occipital Pole |
| | | .0135 | 5.46 | 16 | -78 | -32 | Rt Occipital Pole |
| | 24 | .0031 | 5.91 | 15 | -40 | -40 | Rt Inferior Temporal Gyrus |
| | 81 | .0201 | 5.31 | 27 | -44 | -34 | Rt Medial Temporal Gyrus |
| | 12 | .0231 | 5.27 | -27 | -46 | -33 | Rt Occipital Fusiform Gyrus |
| Visuospatial-early | 124 | .0068 | 5.77 | 40 | -36 | 34 | Rt Supramarginal Gyrus |
| | 69 | .0095 | 5.67 | 32 | -52 | 44 | Rt Superior Parietal Lobule |
| Cognitive | 32 | .0092 | 5.53 | -42 | -45 | 38 | Lt Supramarginal Gyrus |
| | 21 | .0250 | 5.18 | -42 | -48 | -15 | Lt Inferior Temporal Gyrus |
| | 15 | .0187 | 5.28 | -56 | -44 | 22 | Lt Supramarginal Gyrus |
| | 15 | .0200 | 5.26 | -57 | -12 | -15 | Lt Middle Temporal Gyrus |
| | 14 | .0107 | 5.46 | -48 | 4 | 21 | Lt Precentral Gyrus |
| | 14 | .0217 | 5.23 | -44 | -30 | 14 | Lt Parietal Operculum |

Lower scores on the visuo-perceptual-early factor (suggesting worse visuo-perceptual function) were associated with decreased right-hemispheric dominant GM volume predominantly in occipital regions. The largest cluster was located in the cuneal cortex and extended inferiorly into the occipital fusiform gyrus and lingual gyrus. Smaller clusters were also found in the occipital pole, lateral occipital cortex, and medial and inferior temporal gyrus. Occipital and occipito-temporal correlates are consistent with the interpretation of this principal component as capturing early visual and visuoperceptual processes.

Lower scores on the cognitive factor were associated with decreased GM volume across a distributed network of left-hemispheric parietal, temporal and frontal regions. The largest cluster within this network was located in the left supramarginal gyrus. The distributed regions associated with performance on this component have been previously associated with higher cognitive functions relating to attention, working memory and phonological processing, implying higher cognitive demands on the tests loading particularly onto this component.

Lower scores on the visuo-spatial-early factor were associated with decreased GM volume predominantly in the right parietal lobe. The largest cluster was located in the right supramarginal gyrus, and a smaller cluster located in the superior parietal lobule. These correlates are consistent with the interpretation that this component is capturing variation in spatial and visuomotor processes, both of which place demands on the tests loading particularly onto this component.
